# Seed size and its rate of evolution correlate with species diversification across angiosperms

**DOI:** 10.1101/053116

**Authors:** J. Igea, E. F. Miller, A. S. T. Papadopulos, A. J. Tanentzap

**Affiliations:** Department of Plant Sciences, University of Cambridge, Downing St, Cambridge, CB2 3EA, UK; Department of Zoology, University of Cambridge, Downing St, Cambridge, CB2 3EJ, UK; Jodrell Laboratory, Royal Botanic Gardens, Kew, Richmond, TW9 3AB, UK.

## Abstract

Species diversity varies greatly across the different taxonomic groups that comprise the Tree of Life (ToL). This imbalance is particularly conspicuous within angiosperms, but is largely unexplained. Seed mass is one trait that may help clarify why some lineages diversify more than others because it confers adaptation to different environments, which can subsequently influence speciation and extinction. The rate at which seed mass changes across the angiosperm phylogeny may also be linked to diversification by increasing reproductive isolation and allowing access to novel ecological niches. However, the magnitude and direction of the association between seed mass and diversification has not been assessed across the angiosperm phylogeny. Here, we show that absolute seed size and the rate of change in seed size are both associated with variation in diversification rates. Based on the largest available angiosperm phylogenetic tree, we found that smaller-seeded plants had higher rates of diversification, possibly due to improved colonisation potential. The rate of phenotypic change in seed size was also strongly positively correlated with speciation rates, providing rare, large-scale evidence that rapid morphological change is associated with species divergence. Our study now reveals that variation in morphological traits and, importantly, the rate at which they evolve can contribute to explaining the extremely uneven distribution of diversity across the ToL.

## Introduction

Angiosperms are one of the most species-rich clades on Earth and have dominated terrestrial plant communities since the Late Cretaceous [1]. The astounding diversity of flowering plants is distributed extremely unevenly across the ToL. Each of the five most species-rich angiosperm families contains >10,000 species while more than 200 families contain <100 species each [2]. An enduring pursuit in evolutionary biology is to explain this uneven distribution of biodiversity, not only in angiosperms, but also across the whole ToL [3].

Biological traits offer one way to explain disparity in species diversification if they confer adaptation to different environments. Seed mass is one such trait that is particularly important for angiosperms because it integrates across many characteristics of an individual’s life history strategy [4]. Along with adult plant size, seed mass affects survival, reproductive lifespan and dispersal [5]. These life history characteristics contribute to fitness and adaptation, which are the ultimate determinants of whether lineages diversify or go extinct [6]. In support of this idea, seed mass has been shown to correlate negatively with diversification in the Polygonaceae [7], but this has not been investigated across large taxonomic scales. As seed mass varies over ten orders of magnitude in angiosperms, from the minute 1 μg seeds of some orchids to the >18 kg seeds of the sea coconut (*Lodoicea maldivica*), this huge variation may coincide with variation in species diversity. Generalising the direction and magnitude of a link between seed mass and diversification across taxonomic scales has, however, proved difficult. Some life history characteristics encapsulated by seed mass are expected to promote speciation or extinction, while others may simultaneously counteract such effects [8].

The rate of change in key biological traits, such as seed size, can be as important in driving macroevolutionary dynamics as the absolute values of the traits themselves [9]. This is because phenotypic divergence may cause reproductive isolation that results in speciation [10]. Nevertheless, few empirical studies have detected a correlation between rates of phenotypic evolution and lineage diversification ([9,11,12] but see [13]). A correlation between the two may be expected where a trait can change more rapidly in some species than others in response to selective pressures (i.e. high “evolvability” [14]). This rapid change may enable greater access to new ecological niches or quicker establishment of reproductive isolation, thereby increasing the rate of speciation (λ) [15]. In the case of seed mass, the ability to switch rapidly from small seeds with high dispersal ability to larger seeds with lower dispersal ability might promote cycles of rapid colonisation and isolation or permit adaptation to new dispersal vectors in novel environments. Rapid evolution of new phenotypes may also allow individuals to escape harsh environmental conditions and competitive interactions [16], thereby decreasing extinction rates (μ). The overall outcome of these processes on net diversification (r = λ-μ) will ultimately depend upon which of these rates responds more strongly to phenotypic change.

Here, we show that both seed mass and its phenotypic rate of evolution correlate with speciation and extinction across the angiosperm ToL. Our approach combined the most comprehensive phylogenetic timetree available [17] with an unparalleled dataset of seed mass measurements from over 30,000 angiosperm species [18]. We estimated rates of speciation, extinction and seed size evolution across the phylogeny using Bayesian, method-of-moments, and maximum likelihood analyses – each with different assumptions regarding rate variation through time. We then tested whether there were any links between rates of diversification and both seed size and its rate of evolution. Additionally, we examined whether these links were consistent across different methodologies and timescales.

## Results

Our results point to a strong association between angiosperm diversification and rates of seed size evolution irrespective of analytical method or timescale, with weaker evidence for a link between macroevolutionary dynamics and absolute seed size. In the first instance, we calculated rates of speciation (λ), extinction (μ) and seed size evolution using Bayesian Analysis of Macroevolutionary Mixtures (BAMM) [19]. BAMM models rate heterogeneity through time and lineages, and accounts for incomplete taxon sampling. We used a phylogenetic tree that contained 29,703 angiosperm species for the speciation/extinction analysis. The tree was subset to 13,577 species with seed size data for phenotypic evolution analysis. As expected, given the high degree of taxonomic imbalance observed in the angiosperm phylogeny, we found strong support for more than 500 shifts in the rates of diversification. There was also marked heterogeneity in the rates of seed size evolution (Fig. 1), which varied over three orders of magnitude (Fig. S1).

**Figure 1.**
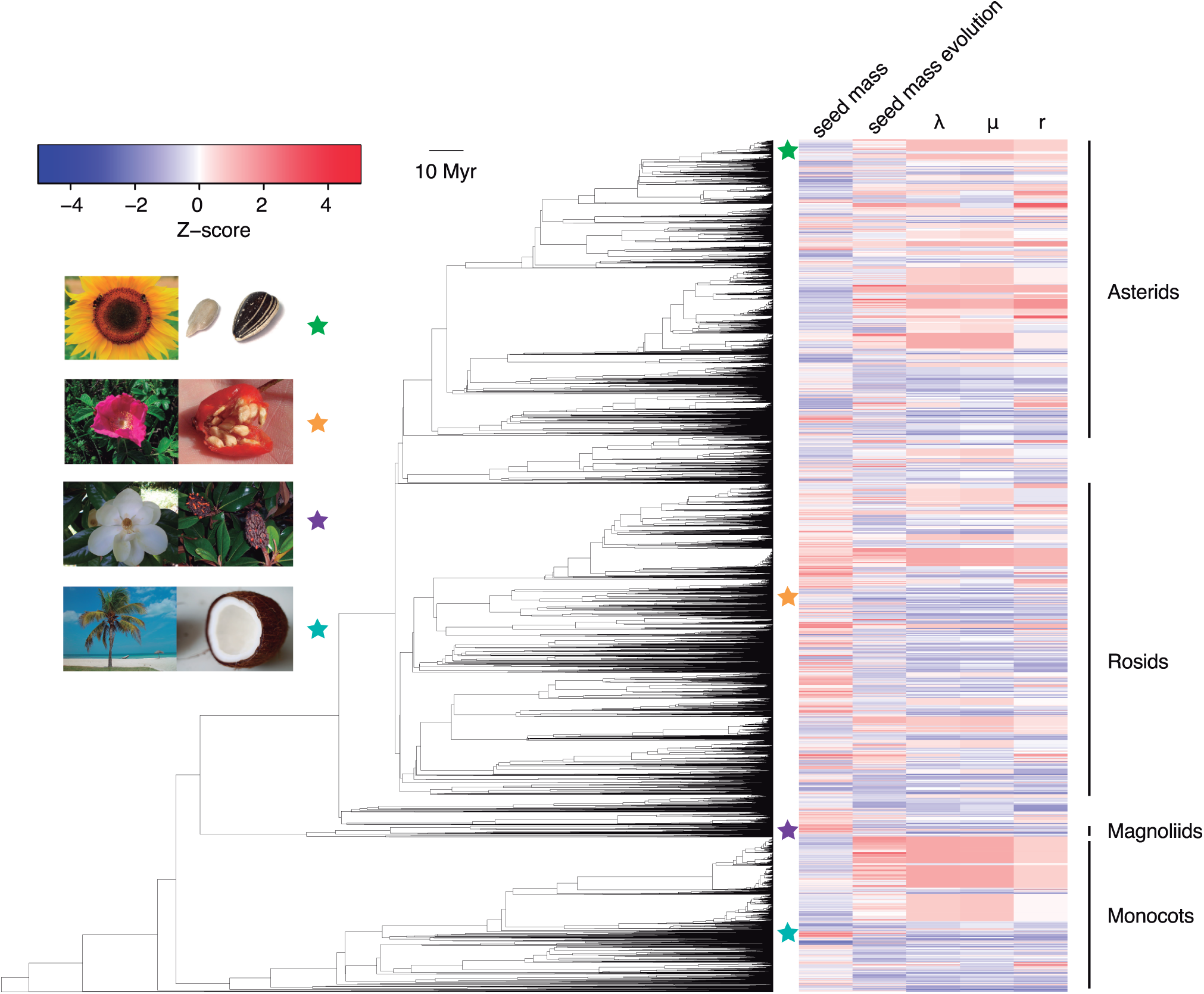
Seed mass and macroevolutionary rates estimated with BAMM across the angiosperm tree of life. Phylogenetic tree of 13,577 species of flowering plants with seed mass, rate of seed mass change, and speciation (λ), extinction (μ) and net diversification (r) rates estimated by BAMM. Seed mass and rate data were standardised to Z-scores so that variation could be directly compared. λ, μ and r were calculated with a larger, 29,703-species tree.

We then estimated whether shifts in macroevolutionary dynamics (λ, μ and r) estimated with BAMM were significantly correlated with absolute seed size and tip-specific rates of seed size evolution by comparing the empirical correlations to a null distribution generated using STructured Rate Permutations on Phylogenies (STRAPP), which is robust to phylogenetic pseudoreplication (see Methods for details) [20]. We were able to link major differences in diversity across angiosperm clades with both the present rate of phenotypic evolution and the absolute value of trait itself. Specifically, increased speciation was associated with a faster rate of seed size evolution (Spearman’s *ρ* = 0.55, p-value < 0.0001; Fig. 2a). Increased extinction rates were similarly associated with higher evolvability (*ρ* = 0.44, p-value < 0.0001; Fig 2b), but given the weaker effect, the net outcome of λ-μ was that diversification rates were positively correlated with phenotypic change (*ρ* = 0.49, p-value < 0.0001; Fig. 2c). We also identified an association between seed size and both speciation (*ρ* = -0.17, p-value = 0.003; Fig. 2d), and extinction rates (*ρ* = -0.17, p-value = 0.003, Fig. 2e). As the correlations with speciation and extinction were in the same direction and of comparable magnitude, and estimates of extinction rates were relatively variable (Fig. 2e), net diversification rates did not change with seed size (*ρ =* -0.12, p-value = 0.077; Fig. 2f). Generally, the observed correlations arose from many phenotypically fast-evolving clades distributed across the phylogeny (Fig. S1) and were robust to prior choice in the BAMM analyses (Fig. S2).

**Figure 2.**
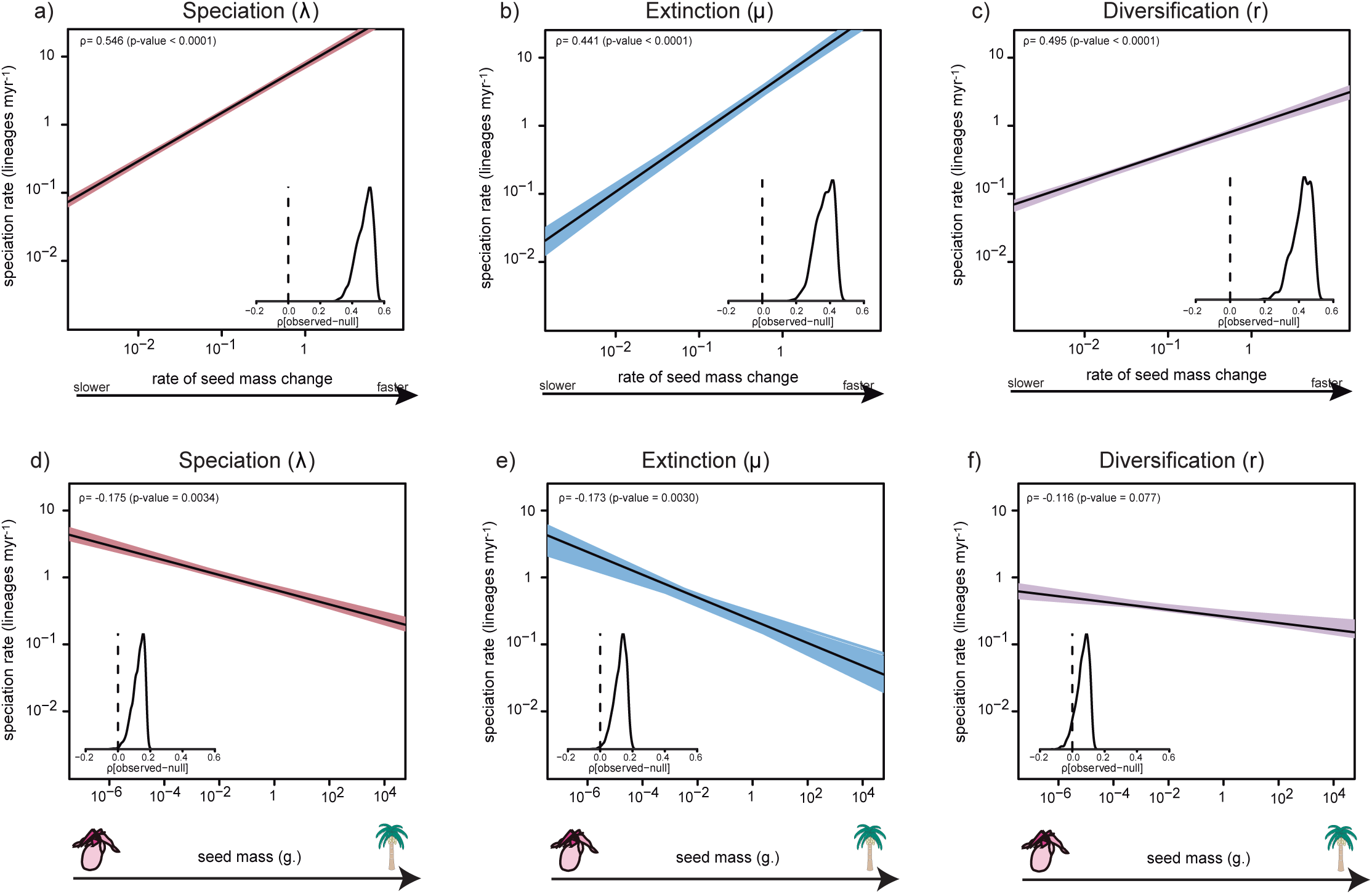
Seed mass and seed mass rate of evolution correlate with macroevolutionary dynamics estimated with BAMM. Spearman correlations were calculated between speciation (λ), extinction (μ), and net diversification (r) and each of a) present-day rate of seed mass change and b) seed mass. Coloured lines are correlations for one sample of the BAMM posterior distribution, bold line is the median. The insets show the density plots of the absolute difference between the observed and null correlation calculated across 1,000 structured permutations of the evolutionary rates on the phylogenetic tree (myr = million years).

Given recent discussion on the reliability of BAMM for estimating diversification rates ([21], but see [22]), we tested the robustness of our results by using alternative methodologies to infer macroevolutionary dynamics across clades at different timescales. Ten, 2 million year-wide time slices from the present up to 20 million years ago were defined. These time slices were used to identify the most inclusive monophyletic clades of ≥4 species in which we had estimated a ≥70% probability of recovering the correct crown age node of the clade (see Methods). For each resulting clade in each time slice, we calculated diversification rates using a method-of-moments estimator, which assumes rates are constant over time [23]. We also fitted a series of time-dependent diversification models to each clade with RPANDA, which uses a maximum likelihood approach to estimate speciation and extinction and allows for incomplete taxon sampling [24] (see Table S1 for a summary of the best fitting models for each time slice). Rates of seed size evolution were estimated within each clade that also had ≥4 species with seed size data by fitting both Brownian motion (BM) and early burst (EB) or accelerating decelerating [25] models of trait evolution. Mirroring the BAMM results, we found a positive correlation between the rate of seed size evolution and speciation rates that was consistent across time slices (Fig. 3a, Fig. S3a). As expected given the weaker association between seed size and speciation found in our BAMM analyses, correlations were generally weaker and non-significant (Fig. 3b), except for one of the time slices (Fig. S3b).

**Figure 3.**
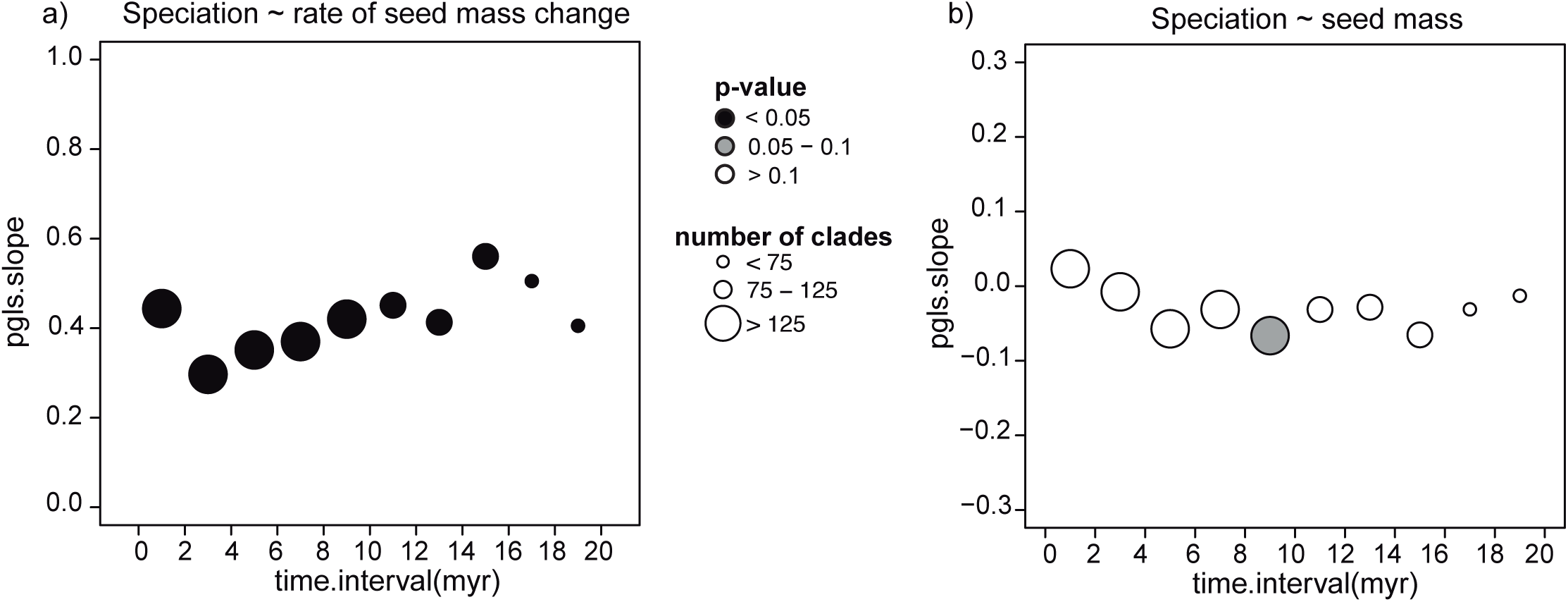
Seed mass and seed mass rate of evolution correlate with speciation in the clade-based analysis. Correlation of (a) rate of seed mass evolution and (b) seed mass with speciation rate (λ) estimated using RPANDA in the clade-based analysis. The strength of correlations is shown as PGLS slopes and were calculated using mean clade-level seed mass across 10 time slices in our species-level phylogenetic tree. The size of the circles represents the number of clades in each time slice plotted at the median age of the time slice. Colour indicates the significance of the slope. A detailed representation of the results in each time slice is given in figures S13 and S14. Correlations calculated with speciation rates obtained with the method-of-moments estimator are given in Fig. S3.

We also found limited evidence that other traits that co-vary with seed size (Tables S2, S3) better explained our results. If our results were explained by seed size being a proxy of some other phenotypic trait that ultimately influenced speciation, we would expect this other phenotypic trait to be strongly correlated with seed size. We would also expect a significantly stronger correlation of speciation with both this trait and its rate of evolution when compared to seed size and its rate of evolution. By comparing the effects of genome size, life cycle, plant height and woodiness across a subset of 1,007 species in our dataset, we found that only the distinction between woody and herbaceous and annual and perennial species were more strongly correlated with macroevolutionary dynamics than absolute seed size. The rate of phenotypic evolution of all the continuous traits (genome size, seed size and plant height) was strongly and similarly correlated with speciation (Fig. S4, Table S4). Importantly, however, neither the rate of evolution nor the absolute values of both genome size and plant height were more strongly associated with speciation than seed size or its rate of evolution (Fig. S4). These results therefore suggest that the correlation between macroevolutionary dynamics and both seed size and its rate of evolution is not simply mediated by other phenotypic traits.

## Discussion

Our study supports the ideas that variation in seed mass and, particularly, its rate of evolution can help explain disparity in diversification across the angiosperm phylogeny by playing a central role in plant life history. As we show with our clade-based analysis, our results are repeatable across many methodologies, varying timescales and are not restricted to a particular taxonomic rank (Figs 2, 3).

The robust association of high rates of phenotypic change with lineage diversification has recently been observed in other taxonomic groups [9,26], but never across the whole of the angiosperm ToL as we find here. Accelerated morphological evolution may allow radiating lineages to occupy more complex adaptive landscapes [27]. Species with greater rate of change in their seed mass (i.e., higher evolvability) could also shift between adaptive peaks or develop reproductive barriers more rapidly. Alternatively, the theory of punctuated equilibria [28], whereby morphological changes can arise from the speciation process, might also explain the connection of phenotypic evolution with species divergence. However, current methods do not allow us to distinguish whether speciation is responding to morphological change or vice versa when reconstructing 250 million years of evolutionary history [9].

Seed mass itself will also co-vary with dispersal ability and environmental tolerance in ways that can change speciation. For example, we found that smaller-seeded genera had faster speciation rates. This may be because smaller-seeded genera generally disperse over larger distances [29], which can promote speciation by creating isolated populations [30]. However, the relationship between dispersal and speciation is highly context dependent. The permeability of a landscape to dispersal determines the dispersal distances that may promote species divergence. For example, long distance dispersal may be needed for isolation to occur in continuous habitats but be less effective in highly fragmented landscapes [31]. Therefore, the weak correlation that we observed between seed size and diversification might reflect contradicting patterns operating in different angiosperm clades. Dispersal syndromes may also modify the effect of seed size on speciation. For instance, species with larger seeds are generally associated with biotic dispersal that distributes seeds over greater distances than wind or gravity dispersal [5]. However, broad-scale predictions for the effects of dispersal syndromes on diversification may be inaccurate, since the former depend on landscape connectivity [8] and can sometimes be inconsistent, e.g. a wind-dispersed seed might be transported by an animal. Detailed contextual data will be necessary to expand upon the mechanisms underlying our findings in specific regions and clades.

Although seed mass is associated with other traits that can affect diversification, there is little evidence that these better explain our observed correlations or that seed size is a mere proxy for one of these other traits. For example, genome size positively correlates with seed mass [32], and faster rates of genome size evolution have been linked to increased speciation in angiosperms [12]. Shorter, smaller-seeded plants also tend to have faster life cycles, which may accelerate mutation rates [33,34] and promote diversification [35]. But unlike other traits [12], both absolute seed size as well as its rate of change were correlated with speciation. Thus, although other traits surely influence diversification [36], we argue that our results generally reflect the role of seed size as a trait that integrates across multiple aspects of life history characteristics in ways that can predictably influence plant macroevolutionary dynamics (Fig. S5).

Our analyses build upon the largest available phylogenetic timetree for angiosperms in ways that do not consider topological and branch length uncertainty. Similar to other megaphylogenies, the low sampling fraction (approximately 10% of described flowering plants) and limited number of phylogenetic markers (a maximum of seven), which were employed in constructing our phylogeny [17], may affect the inference of macroevolutionary estimates [37,38]. In any case, we believe our results, particularly the strong correlation between the rate of seed size evolution and speciation rates may reflect general patterns on how biodiversity is generated across angiosperms. Detailed studies in well-sampled clades can expand upon our findings and reveal different relationships operating in particular groups of organisms.

The approach applied here can help to unravel the processes responsible for generating large-scale asymmetries in biodiversity. It also offers the potential to test how widely-varying traits and their rate of morphological evolution influence other aspects of the evolution and adaptation of flowering plants (e.g. [17]). Clade-specific exceptions arising from local interactions with non-focal traits [39] and specific spatio-temporal contexts will undoubtedly interact with broad-scale macroevolutionary patterns and may modulate the effects of seed mass on diversification. Regardless, our results show that seed size and its rate of evolution correlate with speciation and extinction across the flowering plants. This finding may help to explain why some clades are much more species-rich than others and points to the role of rapid morphological evolution in generating greater levels of diversity.

## Materials and methods

### Seed mass and phylogenetic dataset

Seed mass data for 31,932 species were obtained from the Royal Botanic Gardens Kew Seed Information Database [18]. Species names were standardised with The Plant List (TPL) nomenclature [2] and cleaned using the *Taxonstand* R package [40]. Further processing was performed with the *taxonlookup* R package [41], which is a complete genus-family-order mapping for vascular plants that draws from TPL, the Angiosperm Phylogeny website [42] and a higher-level manually-curated taxonomic lookup [17].

We used the most comprehensive phylogenetic tree for land plants [17,43] that comprises 31,389 species. Taxonomic information for our phylogenetic tree was run through *Taxonstand* and *taxonlookup* as described above to prune it down to angiosperms. The final phylogenetic tree contained 29,703 angiosperm species belonging to 353 plant families (following APG IV, Fig.S6).

### Diversification and phenotypic evolution analyses

Speciation, extinction and net diversification rates were estimated across the 29,703 species tree using BAMM version 2.5.0 [19]. BAMM models shifts in macroevolutionary regimes across a phylogenetic tree using reversible jump Markov chain Monte Carlo (rjMCMC) sampling. The size of the tree precluded a single analysis from readily converging. Therefore, we divided our initial tree into clades of ≥6000 species. This resulted in six monophyletic clades and one additional clade that contained the backbone of the tree (i.e., one representative of the six monophyletic clades) plus the remaining, unassigned species (Fig. S7). We then ran BAMM speciation/extinction analyses for each of the seven clades (six monophyletic clades plus the backbone set). Initial prior settings were calculated with the *setBAMMpriors* function in *BAMMtools* [44], and the *expectedNumberOfShifts* parameter was set at 50. Following [12], we incorporated non-random incomplete sampling information by calculating the proportion of species sampled inside each family and estimated the backbone sampling as the overall proportion of sampled species. *Taxonlookup* was used as a reference for these calculations. Analyses were run for 150 million generations and convergence was verified by plotting chain traces and ensuring that the effective sample sizes of all relevant parameters exceeded 200. The first 100 million generations were discarded as burn-in. The resulting event files for all seven analyses were combined into a single event file, effectively generating one BAMM result set that was then analysed following standard procedure. This clade-level analysis allowed for diversification rate estimation with the most complete dataset that was available. The resulting rate estimates results were strongly correlated with estimates from a single speciation/extinction analysis using the 13,577 species that were present in our seed size database (Fig. S8; r = 0.92, p-value < 0.001).

Rates of seed size evolution were also estimated with BAMM using the phylogenetic tree of the 13,577 species that were present in our seed size database. Initial priors were calculated as above and analyses were run for 300 million generations. The initial 30 million generations were discarded as burn-in. We analysed BAMM prior sensitivity following recent concerns ([45], but see [22]) by re-running both the diversification and the trait evolution analyses for the 13,577 species dataset with different settings for the *expectedNumberOfShifts* parameter of either 25, 50, 100 or 250. These analyses confirmed a low prior sensitivity for the posterior of the number of expected rate shifts (Fig. S2).

Finally, we obtained clade-based measures of diversification and seed size evolution across the species-level tree. Clades were defined as non-overlapping monophyletic groupings of four or more species and their ages were defined using a series of 2-million year wide time slices from the present up to 20 myr ago (see Fig. S13 for the number of clades in each time slice). For each clade, we estimated its sampling fraction by weighting the genus-specific sampling fractions (i.e., the number of congeneric species in the 13,577 species tree divided by the total number of described species for that genus) with the number of species from each genus present in the clade. A minimum clade-specific sampling fraction of 0.3 was used for inclusion in our analyses, ensuring that the crown sampling probability for each clade was at least 0.7 (the actual median clade crown sampling probabilities for the time slices ranged between 0.90 and 0.96). Crown ages for the selected clades were then used to estimate net diversification rates using the method-of-moments estimator [23]. Following standard practice [46,47], we assumed three values of relative extinction fraction, ε = 0, 0.5 and 0.9. Different values did not affect our conclusions (results not shown); therefore we present the results of the intermediate extinction fraction (ε = 0.5). We also used RPANDA to fit a series diversification models that estimated time-dependent rates for each clade. We fitted six different models of diversification: (i) pure birth model with constant λ (speciation rate); (ii) pure birth model with exponential λ; (iii) birth-death model with constant λ and μ (extinction) μ; (iv) birth-death model with exponential λ and constant μ; (v) birth-death model with constant λ and exponential μ; and (vi) birth-death model with exponential λ and exponential μ. We then used AIC-based model selection to select the best fitting model and obtain the corresponding macroevolutionary parameters. Finally, we estimated the rates of seed size evolution by fitting BM and EB models of evolution to the seed size data within each clade using *fitContinuous* from the *geiger* R package [48]. We performed AIC-based model selection to find the best fitting model of trait evolution. More than 99.7% of the clades showed a higher support for the BM model. Additionally, to account for possible biases when analysing clades with many non-congeneric species, we confirmed the results of our clade-based analysis by considering clades that only contained congeneric species (Fig. S9).

### Correlation of diversification and trait evolution

All rate variables were log-transformed for the correlation analyses. Following [20] we treated seed mass evolutionary rates at the tips of the tree as character states and used STRAPP to test for multiple associations between these present-day rates and BAMM-estimated diversification dynamics. We similarly analysed the correlation of absolute seed mass and speciation/extinction dynamics. STRAPP compares the correlation between a focal trait and a macroevolutionary parameter (λ, μ or r) to a null distribution of correlations. The null correlations are generated by permuting the evolutionary rates in the tips of the phylogenetic tree while maintaining the location of rate shift events in the phylogeny. In each case, we calculated the absolute difference between the observed correlation of the macroevolutionary rate and the trait state and the null correlation obtained by the structured permutations across 5000 samples from the BAMM posterior (for an example of the observed correlation in one of the samples in the posterior, see Fig. S10). The reported p-value was the proportion of replicates where the null correlation coefficient was greater than the observed correlation. We found a low type I error associated with our STRAPP correlation analysis (p-value = 0.094, Fig. S11). We also investigated the correlation between the mean speciation and mean phenotypic rates across all the branches in the tree using an ordinary least squares regression (Fig. S12). We note however, that this regression does not correct for phylogenetic dependence in the branch estimates, despite suggesting something about patterns across the whole evolutionary history of the angiosperms.

For the clade-based analyses, we estimated the correlation between speciation rates (measured as λ at present time for the RPANDA analyses) and each of seed size rate of evolution and phylogenetically-corrected mean seed size (i.e., trait value at the root node of the clade under a BM). Correlations were estimated using phylogenetic generalised least square (PGLS) as implemented in the R package *caper* [49] (Fig. S13, Fig. S14). Similar results as presented in the main text were obtained when analysing net diversification instead of speciation rates (Fig. S15). Finally, we similarly analysed the correlation between speciation and each of seed size and its rate of change when selecting only clades consisting of congeneric species. Again, this analysis resulted in a similar pattern as the one presented in the main text (Fig. S9).

### Diversification dynamics and other phenotypic traits

Seed mass is central to a network of inter-correlated traits associated with plant life history that can impact diversification. Some of these traits are genome size or plant C-value (measured as picograms of DNA per haploid nucleus), plant height, life cycle and woodiness. We compared the correlation between macroevolutionary parameters (λ, μ, r) and each of seed mass, seed mass rate of evolution, C-value (i.e., genome size), plant height, life cycle and woodiness across a dataset of 1,007 angiosperm species for which all phenotypic traits could be assembled. Genome content and life cycle data were downloaded from the Plant DNA C-values database [50]. Woodiness data were obtained from [17,51]. Plant height data was obtained from the TRY database [52]. The rates of phenotypic evolution for the continuous traits (seed size, plant height and C-value) were calculated as the phenotypic rate inferred at the tips of the tree in our main BAMM analysis (see above). Surprisingly, mean seed mass did not differ between the 214 strictly annual and 793 perennial plants when accounting for phylogenetic relationships using a phylogenetic ANOVA (Fig. S16, phylANOVA: p-value = 0.308, significance assessed with 1,000 random simulations with *phytools* [53]). In this reduced dataset, we ran STRAPP correlations for each focal trait with the diversification parameters calculated from our main BAMM analysis. We then calculated the absolute differences in the observed and the null correlations between the macroevolutionary parameters and seed mass, C-value, “annuality” (a binary variable specifying whether the species was strictly annual or not), plant height, woodiness (a binary variable specifying whether the species was herbaceous or woody) and the rates of seed size evolution, genome size evolution and plant height evolution. As expected, all phenotypic traits and their rates of evolution were correlated with each other (Fig S5).

### Code availability

Scripts used to carry out the analysis described in the paper and generate the figures will be deposited in Github (https://github.com/javierigea/XXXX) upon acceptance.

## Acknowledgments

We thank V. Soria-Carrasco for help with analyses and D. Rabosky for useful advice on the BAMM analysis. D. A. Coomes, A. J. Helmstetter, T. Jucker and W. G. Lee kindly commented on an earlier draft. J.I. and A.J.T. thank the Gatsby Charitable Foundation, Wellcome Trust and Newton Trust for funding. E.F.M was funded by the BBSRC DTP at the University of Cambridge.

### Author contributions

J.I and A.S.T.P conceived the study. J.I. and E.F.M. performed the analysis. J.I. and A.J.T interpreted the analysis and wrote the manuscript. All authors edited the manuscript.

## Supporting Information captions

**Fig. S1.** Phylogenetic tree of 13,577 angiosperm species with branch colours indicating the rate of seed mass evolution estimated with BAMM. Branches were scaled by speciation rate as determined by a BAMM analysis on a larger 19,703 tree.

**Fig. S2.** Prior and posterior distribution of the number of rate shifts in BAMM for a) the speciation/extinction and b) phenotypic evolution analyses for *expectedNumberOfShifts* = 25, 50 and 100 and 250. The analyses in the main text were carried out with *expectedNumberOfShifts* = 50 for both speciation/extinction and phenotypic evolution analyses.

**Fig. S3.** Seed mass and its rate of evolution are associated with speciation in the clade-based analyses. (a) PGLS slope of the relationship between speciation rate (λ) from the method-of-moments estimator and the rate of seed mass evolution across 10 time slices. Circles are scaled to the number of clades in each time slice while colour indicates the significance of the slope. (b) PGLS slope of the relationship between speciation rate and mean clade seed mass. For a detailed representation of the results in each time slice, see figure S14 and S15.

**Fig. S4.** STRAPP correlations of diversification and phenotypic traits for 1,007 angiosperm species. The distribution of the absolute difference in the observed correlation minus the null correlation is plotted for each trait. The coloured dotted lines indicate the mean of that distribution, and the black dotted line indicates 0; a distribution with mean = 0 would show no association between a focal trait and macroevolutionary dynamics. STRAPP correlation of seed size (shown in red), C-value (shown in yellow), life cycle (shown in light green), woodiness (shown in dark green), height (shown in blue), seed size rate (shown in dark blue), C-value rate (shown in purple); and height rate (shown in pink) with a) speciation rate (λ), b) extinction rate (μ), and c) net diversification rate (r).

**Fig. S5.** Proposed effects of seed mass and other life history traits on diversification (solid lines). Dashed lines indicate correlations between life history traits. Numbers indicate reference where the link is proposed.

**Fig. S6.** Phylogenetic tree of 353 angiosperm families with representatives in our analyses. The red bars indicate the levels of sampling for each family.

**Figure S7.** Angiosperm phylogenetic tree collapsed to monophyletic clades of ≤6000 species. The name of one representative species per clade is shown, and the numbers in parentheses indicate the number of species included in each clade. The BAMM analyses were carried out for six monophyletic clades (shown in red, yellow, green, blue, dark blue and pink) and one “backbone” analysis with the remaining clades (shown in grey) and one representative of each of the six monophyletic clades.

**Fig. S8.** Comparison of the speciation rates at the tip of the tree obtained with the complete Zanne tree (29,703 species) and the seed size filtered tree (13,577 species). The dotted line represents the 1:1 reference line.

**Figure S9.** Correlation of speciation with seed mass and seed mass rate of evolution in the clade-based analysis only considering congeneric species. (a) PGLS slope of the relationship of speciation rate - estimated with the method-of-moments estimator - with mean clade seed mass across 10 time slices. The size of the circles represents the number of clades in each time slice while the colour indicates the significance of the slope. (b) PGLS slope of the relationship of speciation rate and the rate of seed mass evolution.

**Fig. S10.** Correlation between speciation rate and rate of seed size evolution in a random sample of the BAMM posterior. The dotted line represents the Spearman correlation (ρ = 0.47, p-value < 0.001)

**Fig. S11.** Type I error analysis. We estimated the type I error rate of our analysis by simulating neutral traits on the angiosperm phylogenetic tree. We performed 1,000 simulations and then ran 1,000 STRAPP tests with each simulated dataset. We estimated the corresponding p-values for the association between traits and diversification and calculated the type I error as the proportion of datasets where a significant association (p-value < 0.05) was detected.

**Fig. S12.** Correlation of mean speciation and mean phenotypic rates across all the branches of the angiosperm phylogenetic tree. The dotted line is the ordinary least squares regression (R^2^ = 0.31, p-value < 0.001)

**Fig. S13.** Correlations between clade rate of seed size evolution and speciation rate(estimated with RPANDA) across time slices from: (a) 0 to 2 million years (myr); (b) 2 to 4 myr; (c) 4 to 6 myr; (d) 6 to 8 myr; (e) 8 to 10 myr; (f) 10 to 12 myr; (g) 12 to 14 myr; (h) 14 to 16 myr; (i) 16 to 18 myr; and (j) 18 to 20 myr. The degrees of freedom (df) are equivalent to the number of clades minus one.

**Fig. S14.** Correlations between mean clade seed mass and speciation rate (estimated with RPANDA) across time slices from: (a) 0 to 2 million years (myr); (b) 2 to 4 myr; (c) 4 to 6 myr; (d) 6 to 8 myr; (e) 8 to 10 myr; (f) 10 to 12 myr; (g) 12 to 14 myr; (h) 14 to 16 myr; (i) 16 to 18 myr; and (j) 18 to 20 myr. The degrees of freedom (df) are equivalent to the number of clades minus one.

**Fig. S15.** Correlation of (a) rate of seed mass evolution and (b) seed mass with net diversification rate (r) estimated using RPANDA in the clade-based analysis. The strength of correlations is shown as PGLS slopes and was calculated using mean clade-level seed mass across 10 time slices. The size of the circles represents the number of clades in each time slice while the colour indicates the significance of the slope.

**Fig. S16.** Mean genus seed mass of strict annual (n = 214) and perennial (n = 793) genera. No significant difference between the means of the two groups was found when accounting for phylogeny (phylANOVA: p-value = 0.308, significance assessed with 1,000 random simulations).

**Fig. S17.** Correlations between clade rate of seed size evolution and speciation rate(estimated with the method-of-moments estimator) across time slices from: (a) 0 to 2million years (myr); (b) 2 to 4 myr; (c) 4 to 6 myr; (d) 6 to 8 myr; (e) 8 to 10 myr; (f) 10 to 12 myr; (g) 12 to 14 myr; (h) 14 to 16 myr; (i) 16 to 18 myr; and (j) 18 to 20 myr.

**Fig. S18.** Correlations between mean clade seed mass and speciation rate (estimated with the method-of-moments estimator) across time slices from: (a) 0 to 2 million years (myr); (b) 2 to 4 myr; (c) 4 to 6 myr; (d) 6 to 8 myr; (e) 8 to 10 myr; (f) 10 to 12 myr; (g) 12 to 14 myr; (h) 14 to 16 myr; (i) 16 to 18 myr; and (j) 18 to 20 myr.

**Table S1.** RPANDA diversification models for the clade-based analyses. For each 2-million year (myr) time slice, we counted the number of clades where the best-fitting model was either i) birth-death model with constant λ (speciation) and μ (extinction) (lambda.cst.mu.cst); pure birth model with constant λ (lambda.cst.mu0); pure birth model with exponential λ (lambda.exp.mu.0); birth-death model with exponential λ and constant μ (lambda.exp.mu.cst); birth-death model with exponential λ and exponential μ (lambda.exp.mu.exp); or birth-death model with constant λ and exponential μ (lambda.cst.mu.exp).

**Table S2.** Correlations of seed size and other phenotypic traits. Trait values were obtained from a 1,007 species tree where all species had data for seed size, C-value and plant height. The values are the slopes of the PGLS regressions and asterisks denote statistically significant correlations (p-value < 0.05).

**Table S3.** Correlations of seed size rate of evolution and other phenotypic rates of evolution. Rate values were obtained from a 1,007 species tree where all species had data for seed size, C-value and plant height. The values are the slopes of the PGLS regressions and asterisks denote statistically significant correlations (p-value < 0.05).

**Table S4.** STRAPP correlations (*rho*) for 1,007 species of angiosperms with seed size, genome size (i.e., C-value), life cycle, height, woodiness data and rates of seed size, C-value and height evolution. Significant correlation are shown in bold and p-values are shown in parentheses.

